# A Hydrogel Microneedle-assisted Assay Integrating Aptamer Probes and Fluorescence Detection for Reagentless Biomarker Quantification

**DOI:** 10.1101/2021.10.14.464448

**Authors:** Hanjia Zheng, Amin GhavamiNejad, Peyman GhavamiNejad, Melisa Samarikhalaj, Adria Giacca, Mahla Poudineh

## Abstract

Analyzing interstitial fluid (ISF) via microneedle (MN) devices enables patient health monitoring in a minimally invasive manner and at point-of-care settings. However, most MN-based diagnostic approaches require complicated fabrication processes or post-processing of the extracted ISF. Here we show in-situ and on-needle measurement of target analytes by integrating hydrogel microneedles (HMN) with aptamer probes as the target recognition elements. Fluorescently tagged aptamer probes are chemically attached to the hydrogel matrix while a crosslinked patch is formed. We use the assay for specific and sensitive quantification of glucose concentrations in an animal model of diabetes to track hypoglycemia, euglycemia, and hyperglycemia conditions. The assay can track the rising and falling concentrations of glucose and the extracted measurements closely match those from the gold standard techniques. The assay enables rapid and reagentless target detection and can be readily modified to measure other target analytes *in vivo*. Our system has the potential to improve the quality of life of patients who are in need of close monitoring of biomarkers of health and disease.

---

Transdermal biosensing can bring us one step closer to personalized and precision medicine, as it enables the continuous tracking of patient health conditions in a non- or minimally invasive manner^1,2^. Transdermal biosensors analyze interstitial fluid (ISF), the fluid which is present in the lowermost skin layer of the dermis, for biomarker measurements^2–4^. Compared to other body fluids, ISF has the most similar molecular composition to blood plasma^5^, in addition to possessing other unique features including biomarkers of medical relevance^2,6^. Simple and effective methods that enable the comprehensive analysis of ISF can lead to transformative advances in bio-diagnostic technologies^3,6–8^. These approaches are not only minimally invasive and painless, but also ideally suited for point-of-care and resource-limited settings^1,3,6,9^.

Microneedle (MN)-based techniques have been introduced as effective approaches for simple ISF extraction with the potential of integrating diagnostics^7,8,10,11^. Different types of MNs implement various strategies to obtain ISF, for example, hollow MNs operate based on negative pressure^12,13^; porous MNs use capillary force^13,14^; and the most recent one, hydrogel-based MNs (HMNs) employ material absorption property^3,7^. HMNs with a length less than 1000 μm and tips much sharper than hypodermic needle enable efficient piercing of the stratum corneum (outer layer of the skin) and the formation of microscale ISF extraction channels^4,15,16^. Compared to other MNs, HMNs possess several advantages, including increased and rapid ISF extraction, high biocompatibility, lower fabrication cost, higher production yield, and most importantly ease of insertion and removal without causing skin damage^4,7,16–18^.

Integrating biosensors on MNs enables in-situ ISF characterization^3,4^. Hollow, metallic MN devices combined with enzymes have been implemented for real-time monitoring of various metabolites, electrolytes, and therapeutics^3,7,19^. The main obstacles with hollow MN applications are the complex fabrication protocols and the potential risk of MN clogging^8^. MNs functionalized with antibodies and aptamers - short single-stranded DNA capable of specific binding to a target molecule-have been recently reported for the specific capture of target biomarkers in ISF, followed by *ex vivo* analysis^20,21^. Although these MN devices allow for on-needle biomarker detection in ISF, the sensor still needs post-processing steps, such as washing and adding detection reagents to detect targets of interest.

In this paper, for the first time, we report a fluorescent HMN biosensor based on methacrylated hyaluronic acid (MeHA) for on-needle and reagentless capture and detection of any biomarkers of interest. Our reagentless fluorescence assay for minimally invasive detection (RFMID) integrates a novel, rapid, and simple approach to link aptamer probes to the MeHA matrix. We demonstrate the application of our biosensor for *ex vivo* detection of adenosine triphosphate (ATP) and glucose, where HMN arrays functionalized with aptamer probes can detect the analyte concentrations with high sensitivity and specificity. We also show that the RFMID can be employed for tracking rising and falling levels of glucose in an animal model of diabetes. Specifically, the RFMID can accurately track severe hypoglycemia range, which cannot be detected using the commercially available glucose monitoring devices. The proposed RFMID technique is expected to pave the way for the next generation of real-time, continuous biosensors.

## Results and Discussion

### RFMID detection strategy

The RFMID integrates HMNs with aptamer probes as biorecognition elements to selectively capture the target analyte in a minimally invasive manner. HMNs are fabricated using MeHA, a highly swellable and biocompatible polymer that has been previously employed for enhanced ISF extraction and off-site target detection^4^. MeHA was synthesized by modifying HA with methacrylic anhydride^4,22^. The degree of methacrylation was determined to be 20% by integration of methacrylate proton signals at 6.1, 5.7, and 1.8 ppm to the peak at 1.9 ppm related to the N-acetyl glucosamine of HA^23^ (**Figure S1**). A novel approach was employed to link aptamer probes into MeHA-HMNs by introducing an acrydite group to the proximal end of the aptamer. In the presence of a photoinitiator (PI) and under UV exposure, the acrydite group forms a covalent linkage with MeHA, enabling the attachment of aptamer probes to the HA network (**Figure 1a & 1b**). Additionally, a crosslinked hydrogel patch is formed via covalent attachment of unreacted MeHA carbon-carbon double bonds with the crosslinking agent.

**Figure 1:**
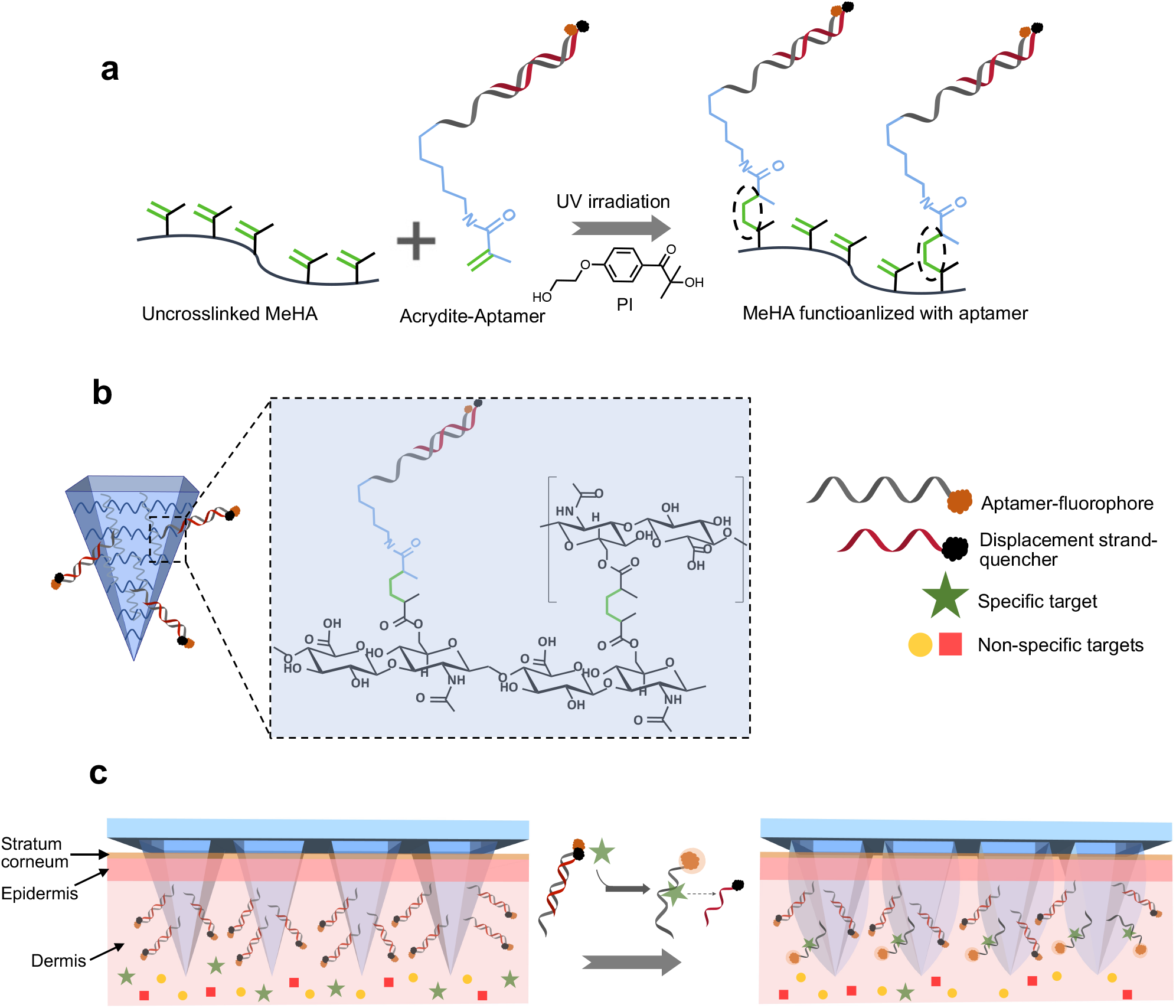
Overview of RFMID functionalization and sensing strategy. **a**, The RFMID employs a covalent binding to attach aptamer probes into MeHA. The Cy3 conjugated aptamer with an acrydite group on the 5’ end is pre-hybridized with the corresponding competitor strand to form the aptamer-quencher complex, which can be covalently attached to the hydrogel matrix during the MeHA crosslinking process and in the presence of PI and under UV exposure. **b**, The RFMID’s needles consist of crosslinked MeHA that is immobilized with aptamer-quencher complex. **c**, The RFMID uses a reagentless process for target detection. Upon insertion, the RFMID penetrates through stratum corneum and epidermis and rapidly swells to extract transdermal ISF. During this process, fluorophore conjugated aptamer probes selectively bind to the specific target, leading to dissociation of the quencher strand and producing fluorescent signal.

For reagentless target detection, we used aptamer probes and a strand displacement strategy^24,25^. Briefly, we hybridized Cy3 fluorophore-conjugated aptamers with a DNA competitor strand that has been conjugated to a quencher (Dabcyl) molecule, called the quencher strand, and coupled the complex to MeHA through covalent linkage. The aptamer-quencher complex retains the fluorophore and quencher in proximity, producing no signal in the absence of a specific target. When the aptamer binds to the target, the quencher competitor strand dissociates and alleviates quenching of the fluorophore, producing a signal without requiring for any post-processing steps such as washing or adding a detection reagent. Despite that fact that the displacement-based analyses often suffer from low sensitivities, this is not the case in our assay because the binding of an aptamer to its target molecule is more thermodynamically stable than the binding to its complementary stand^26^. **Figure 1c** shows a schematic of the RFMID reagentless detection principle. After fabrication, the RFMID is pressed through the skin for ISF extraction and target capture. Because non-specific binding of background chemicals does not cause dissociation of the quencher strand, the probe only responds to the binding of the specific target even when the background concentration is many orders of magnitude higher. This important feature enables our sensor to operate in complex mediums such as ISF without the need for sample preparation or added reagents.

### RFMID fabrication and testing

We fabricated the RFMID using a negative polydimethylsiloxane (PDMS) mold (**Figure 2a, i**). MeHA, PI and the crosslinking agent, N,N’-methylenebisacrylamide (MBA), were dissolved in the corresponding buffer solution and applied to the PDMS mold.

**Figure 2:**
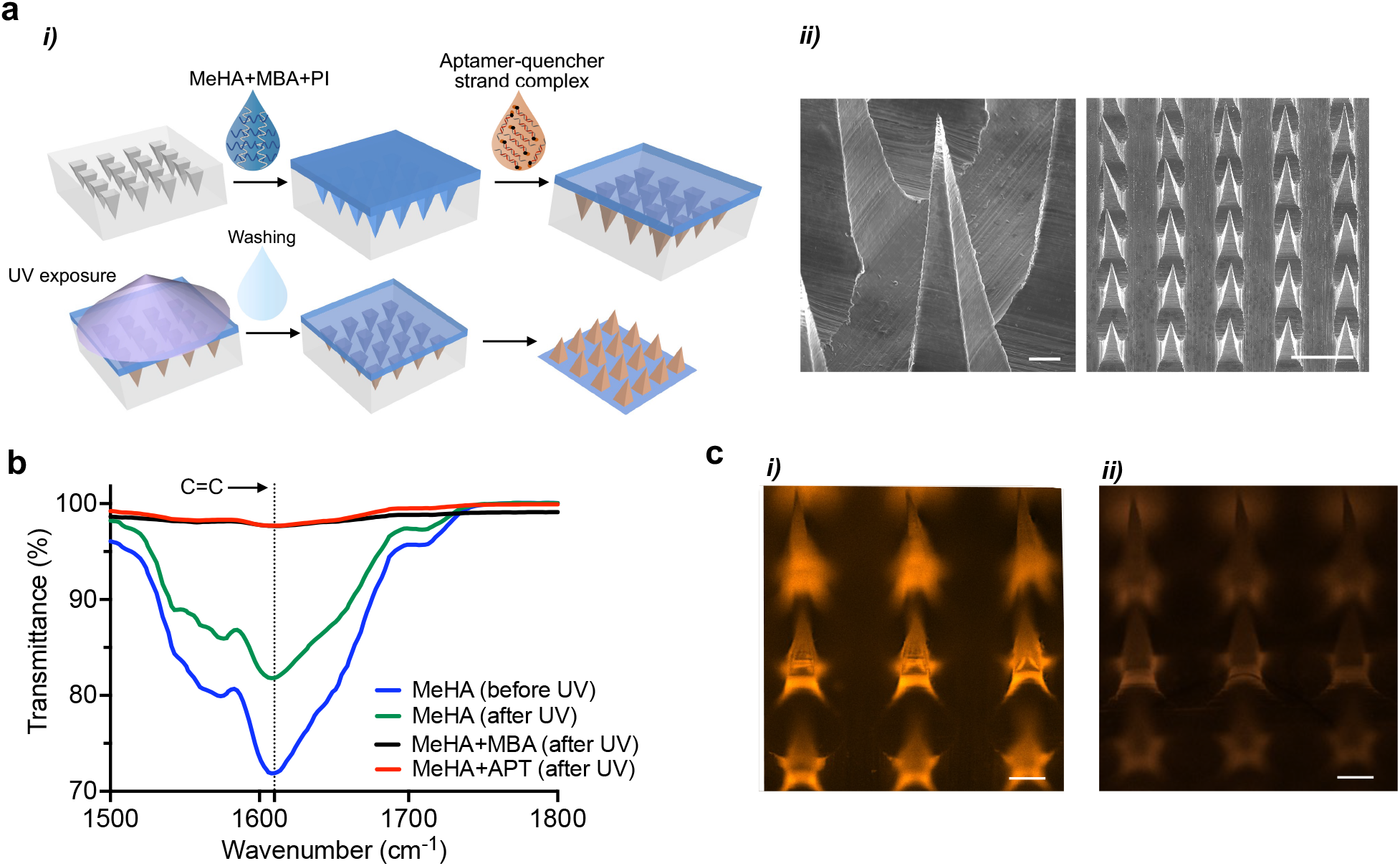
RFMID fabrication and functionalization. **a**, Schematic showing the fabrication process of RMFID: *i)* MeHA solution (40% w/v in Glucose or ATP binding buffer), PI, and MBA are casted into silicon mold, followed by the addition of aptamer-quencher complex. After drying, RFMID is crosslinked for 20 min under UV exposure and then washed to remove unbounded aptamer. RFMID device is then removed from the silicon mold and is ready to be applied for biomarker measurement. *ii)* Scanning electron microscopy (SEM) images showing the morphology of fabricated RFMID from the single needle view (left) and side view (right). Scale bar, 50 μm (left) and 500 μm (right). **b**, FTIR spectra of MeHA thin films before and after UV irradiation and MeHA with MBA or with ATP aptamer probe (APT) after UV irradiation. The reduction in transmittance peaks of hydrocarbyl group at 1633 cm^−1^, for crosslinked samples with aptamer probe or MBA shows the covalent attachment of the aptamer or MBA to the crosslinked patch. **c**, Fluorescence microscopic images of RMFID patches functionalized with an aptamer probe only (left) and aptamer-quencher complex (right). Scale bar, 250 μm.

Before complete drying, the hybridized aptamer-quencher complex was added to the mold and the patches were left to dry. The patches were then exposed to UV for both aptamer linking and patch crosslinking. To remove the unbound aptamer probes, the patches were washed twice inside the mold to avoid the deformation of patch needles. The HMN patches were then removed from the mold and exposed to UV for another 5 min through the needle side to ensure formation of a crosslinked patch (not shown in **Figure 2a**). Following this process, we were able to fabricate HMN patches with sharp needles (**Figure 2a, ii**) that are critical for effective skin penetration^4,20^.

We first investigated the chemical structure of hydrogel biosensor and efficiency of linking the aptamer probes to MeHA using Fourier-transform infrared spectroscopy (FTIR) (**Figure 2b**). The spectrum of MeHA sample before UV exposure showed a strong peaks at 1633 cm^1^ (C=C)^4^, which significantly decrease in intensity after UV irradiation in MeHA sample with MBA, indicating that the carbon-carbon double bonds of MeHA were broken to form the covalent linkage with MBA. This reduction in the peak intensity was found to be minimal in the pure MeHA sample after UV irradiation. However, a similar trend with MeHA+MBA sample was observed for MeHA mixed with aptamer probes conjugated with acrydite group (MeHA+APT) after UV irradiation, suggesting a successful attachment of aptamer probes to the MeHA network. The successful aptamer linkage was also confirmed via fluorescence measurement after the fabrication process. We applied the aptamer-quencher complex or only aptamer probes and observed the patches under fluorescence microscopy. The aptamer-only patch (**Figure 2c, i**) shows an elevated fluorescence signal compared to the patch functionalized with aptamer-quencher complex (**Figure 2c, ii**), demonstrating both the successful aptamer linkage and quenching capabilities of Dabcyl.

We next investigated the effect of aptamer probes on the swelling capability and mechanical strength of the RFMID patches. The swelling capability of the RFMID patches were tested by measuring the patch’s weight before and after application through an agarose hydrogel for 10 mins. We observed that the presence of aptamer probes does not have any significant effect on the swelling ability of MeHA hydrogel network (**Figure 3a**). The slight increase in the swelling might be due to the lower crosslinking density in the RFMID patches (**Figure 1b**). We also observed a reduction of swelling by extending the crosslinking period (**Figure S2**), which agrees with previous work reported on MeHA-HMNs^4^. The mechanical strength of our MeHA-HMN patch was also evaluated through a compression test. HMN patches with and without aptamer show similar load versus displacement profiles (**Figure 3b**), which indicates the mechanical strength is not influenced by the aptamer probes. The compression test result also shows that HMNs in this study can exert more than 0.6 N, which is sufficient for successful piercing through the skin^4^. The capability of the RFMID for target capture or recovery was investigated using glucose as a target molecule via testing three HMN samples; a blank HMN patch with no aptamer probes, a HMN patch functionalized with glucose aptamer probes (Glu apt HMN), and a HMN patch functionalized with glucose aptamer probes where upon target capture the aptamer probes were degraded using ultrasonication (Glu apt-US HMN)^27^ (**Figure 3c**). The three HMN patches were then pressed through an agarose hydrogel loaded with varying concentrations of glucose. The diffused or captured targets by blank HMN and Glu apt HMN were then recovered by centrifugation. For Glu apt-US HMN, prior to centrifugation, the patches were sonicated to degrade the aptamer and release the captured target. As shown in **Figure 3c**, the blank HMN and Glu apt-US HMN show a high recovery rate where the absence and the degradation of aptamer probes, respectively, allow for the release and recovery of the target molecules. While in Glu apt HMN sample, the presence of aptamer probes hinders the full recovery of glucose. This experiment ultimately indicates that the analytes diffuse into the patch needles and are captured by aptamer probes for subsequent detection. To visualize the target recovery capability, HMN patches were pressed through an agarose hydrogel loaded with Rhodamine B (RhoB) then the diffused dye was recovered (**Figure 3d & S3**).

**Figure 3:**
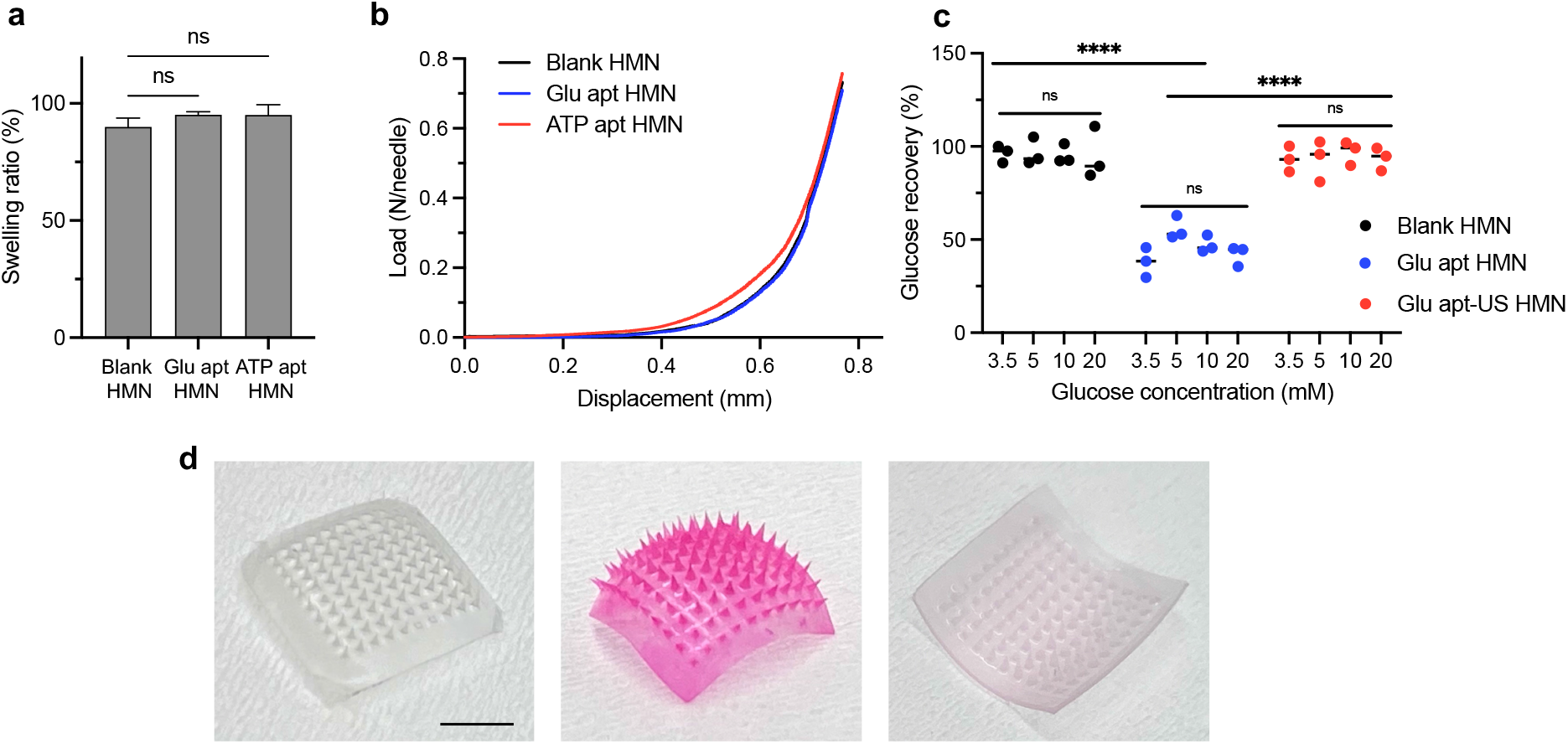
RMFID characterization. **a**, Swelling ratio of a blank HMN patch without aptamer and HMN patches functionalized with glucose or ATP aptamer probes. The presence of aptamer does not affect the swelling capability of HMN patches. Data are mean ± s.d. n = 3 repeated tests per group. The data among Glu apt HMN and ATP apt HMN groups was not significantly different from the blank HMN group (ns, not significant, *P* > 0.9999 by the ordinary one-way analysis of variance (ANOVA) with Turkey’s multiple comparison test). **b**, Mechanical compression test for blank HMN and HMN patches functionalized with glucose and ATP aptamer probe. A compressive force in the longitudinal direction of the HMNs was generated by a moving transducer at a speed of 0.5 mm/min. **c**, A blank HMN patch and HMN patches functionalized with the glucose aptamer probes (Glu apt HMN) were inserted into agarose hydrogels containing various concentrations of glucose (3.5, 5, 10, 20 mM) for 10 min to capture or recover glucose. For one group of aptamer HMNs, upon target capture, the aptamer probes were degraded by sonication (Glu apt-US HMN). The glucose was then recovered by centrifugation at 4,880 rpm for 5 min. Data is expressed as mean ± s.d. n = 3 replications per group. Data in Blank HMN and Glu apt-US HMN groups are significantly higher than Glu apt HMN group (**** *P* > 0.9999 by two-way ANOVA with Geisser-Greenhouse correction). Among each group, data of various glucose concentration group is not significantly different (ns, not significant, *P* > 0.9999 by two-way ANOVA with Geisser-Greenhouse correction). **d**, Optical images of a HMN patch before (left) after (middle) solution extraction from an agarose hydrogel containing 100 mg/mL RhoB, followed by the RhoB recovery (right). Scale bar, 5 mm.

### *In vitro* and *Ex vivo detection of ATP and glucose*

To investigate our sensor’s capability for biomolecule detection, a series of experiments were conducted to measure varying concentrations of glucose (**Figure 4a**) and ATP (**Figure 4b**) *in vitro* using HMNs linked with glucose aptamer-quencher^28,29^ complex and ATP aptamer-quencher^30^ complex, respectively. We first studied the capability of the glucose/ATP aptamer-quencher strand complex for target detection and optimized the ratio of aptamer to quencher strand (**Figure S4**). HMN patches linked with the optimum ratio of aptamer and quencher strand were then applied on agarose hydrogels loaded with different concentrations of ATP or glucose for 10 min. We measured the patch fluorescence intensity before and after application to agarose hydrogel and observed that by increasing the target concentration, the signal intensity increases, demonstrating the target capture and dissociation of the quencher strand. We estimated the limit of detection (LOD) of our biosensor to be three times the standard deviation of the fluorescence signal intensity from a blank agarose hydrogel (See **Methods**). Our HMN sensor achieved a LOD of 2.5 mM for glucose and 0.2 mM for ATP measurements. To further explore the sensitivity and reliability of detection, cross-reactivity among glucose and ATP was examined by applying HMN patches functionalized with glucose (ATP) aptamer into an agarose hydrogel loaded with a definite concentration of glucose (ATP) while the concentrations of ATP (glucose) increased. We observed that for both glucose and ATP, the changes of the fluorescence intensity for the non-specific target were almost unnoticeable, indicating the high selectivity of the sensor (**Figure 4c**). The cross-reactivity of RFMID device for glucose detection was also studied against fructose, uric acid, insulin, 3-β-Hydroxybutyrate-the dominant biomarker of ketone formation^31^ (**Figure S5**). We observed that the addition of common interfering agents (with a concentration higher than their physiological levels) does not affect the RFMID measurement which confirms excellent selectivity of the sensor.

**Figure 4:**
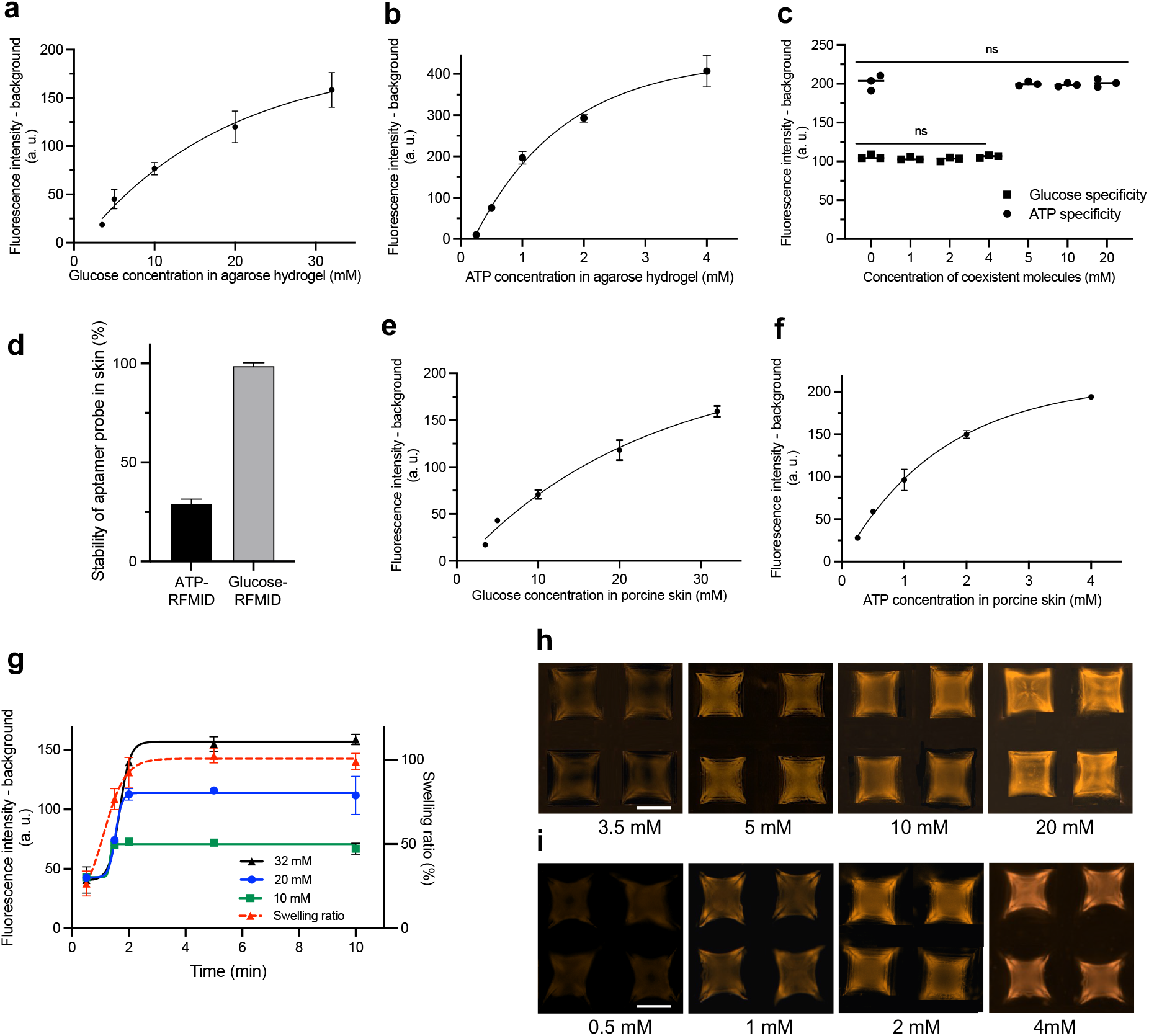
*In vitro* and *ex vivo* detection of ATP and glucose. **a**, The RFMID devices functionalized with glucose or **b**, ATP aptamer-quencher complex were applied into agarose hydrogels containing varying concentrations of glucose (3.5, 5, 10, 20, 32 mM) or ATP for 10 mins (0.25, 0.5, 1, 2, 4 mM). The fluorescence signal intensity was measured before and after applying the patches and the difference in signal intensity was reported. **c**, The specificity of RFMID devices for capturing the specific target was tested. The ATP-RFMID devices were applied into agarose hydrogels containing 1 mM of ATP while glucose concentrations increased (0, 5, 10, 20 mM). Similarly, the glucose-RFMID devices were applied into agarose hydrogel containing 20 mM of glucose while ATP concentrations increased (0, 1, 2, 4 mM). **d**, Stability of ATP and glucose aptamer probes in skin. HMN patches functionalized with only aptamer probes were applied to porcine skin or a blank agarose hydrogel. Upon insertion, the corresponding quencher strands were added to the patches. The reduction of fluorescence signal intensity of the patches applied to skin or agarose were then measured and compared. The graph shows 1-[(fa-fs)/fa] as a measure of aptamer stability, where fa and fs are the reduction in fluorescence signal for HMN patches applied into the agarose hydrogel or skin, respectively, upon addition of quencher strands. The RFMID devices functionalized with **e**, glucose or ATP **f**, aptamer-quencher complex were applied into porcine skin equilibrated with varying concentration of glucose (3.5, 5, 10, 20, 32 mM) or ATP (0.25, 0.5, 1, 2, 4 mM) for 5 min. The fluorescence signal intensity was measured before and after applying the patches and the difference in signal intensity was reported. Since some ATP aptamer probes are degraded in skin (based on experiments in **d**), the level of fluorescence signal is less compared with agarose hydrogel experiment in **b. g**, Glucose-RFMID devices were applied through porcine skin equilibrated with different glucose concentrations (10, 20, and 32 mM) for different durations. The swelling of patches and the fluorescence response were then measured. 2 min was found to be the optimal time of HMN application. **h**, Fluorescent microscopic images of glucose-RFMID and **i**, ATP-RFMID after capturing varying concentrations of ATP (0.5, 1, 2, 4 mM) or glucose (3.5, 5, 10, 20, 32 mM) loaded in the agarose hydrogel. Data are presented as mean ± s.d. a.u., arbitrary units.

Upon successful detection of target analytes in agarose hydrogel, the sensor capability for in situ biomarker measurement was tested using an *ex vivo* skin model. First, the degradation of both ATP and glucose aptamer probes via nucleases present in the skin was examined. To this end, functionalized HMN patches with only aptamer probes were applied through a blank porcine skin or agarose hydrogel (with no nucleases present). We observed that the fluorescence intensity of the patches did not change after insertion. After penetration, the quencher strand was added to the HMN patches and a reduction in fluorescence signal was observed. The reduction of fluorescence signal intensity of the patches applied to skin or agarose were then measured and compared. The fluorescence signal reduction in HMNs penetrated through skin was 98% and 29% of the ones inserted into blank agarose hydrogel for glucose and ATP aptamer probes, respectively, (**Figure 4d**) indicating that most of ATP aptamers became degraded in the skin and could not be hybridized to the corresponded quencher strands. These results confirm that the glucose aptamer has a great stability in skin while the ATP aptamer might not be sufficiently stable to permit target detection in a complex ISF environment.

The RFMID was then employed for glucose (**Figure 4e**) and ATP (**Figure 4f**) detection using porcine skin equilibrated with different concentrations of the target analytes. Microneedle traces were evident in the porcine skin, showing an efficient skin penetration (**Figure S6**). We used these data to construct standard curves that correlate fluorescence signal intensity to glucose or ATP concentration in skin ISF. Our device achieved a LOD of 1.1 mM for glucose and 0.1 mM for ATP measurements in skin ISF. Our glucose detection limit and dynamic range cover the clinical hypoglycemia, euglycemia and hyperglycemic ranges. Our system can potentially be employed to reliably quantify pre- and post-prandial glucose concentrations in patients with diabetes while the accuracy of most of commercially available glucose monitoring devices is still the lowest within the hypoglycemic range^32^.

To determine the shortest timescale for effective capture of target analytes, glucose-RFMID patches were applied on the porcine skin equilibrated with various concentrations of glucose for different durations. 2 min of microneedle patch administration was found to be sufficient to capture and detect glucose (**Figure 4g**) and longer administration (5 and 10 min) did not change the measured fluorescence. The fast response is because of the swelling characteristic of MeHA-HMNs that can reach their maximum swelling within 2 min, enabling increased and rapid ISF extraction and thus target capture. The 2 min response time is also independent of the target concentration and increased glucose concentration does not affect the time needed for the target molecules to diffuse into the patch needles. Previously reported MN-based diagnostic devices require at least 140 min to detect the target analytes (**Table S1**), including the time needed for MN application, washing, and adding the detection agents^20,21,33^. **Figure 4h** and **4i** show that the fluorescence intensity of MN patches increases with the rising glucose and ATP concentration, respectively, confirming elevated dissociation of quencher strand and target capture.

### In vivo glucose detection in animal models of diabetes

Having demonstrated our platform’s ability to detect glucose *in vitro* and *ex vivo* sensitively and accurately, we evaluated its performance *in vivo* using a streptozotocin-induced rat model of diabetes. Prior to the animal experiment, the cytotoxicity of the composite materials was evaluated in NIH-3T3 fibroblast cells using MTT assay (**Figure S7**). Results showed that NIH-3T3 fibroblast viability was not significantly influenced, suggesting that the RFMID materials were biocompatible. RFMID patches were inserted on the rat’s dorsal skin side (**Figure 5a**). HMN patches efficiently penetrated the tissue as evidenced by the microneedle traces and the skin recovered well post-treatment (**Figure 5b**). The HMN punctures disappeared gradually within 15 min of removing the patch. This is due to the biocompatibility and minimal invasiveness of the RFMID patches. The diabetic rats were fasted for 5 hours prior to the experiment and treated first with 4 IU kg^−1^ dose of human recombinant insulin subcutaneously to lower blood glucose and after hypoglycemia was reached with 30% glucose intraperitoneally to increase blood glucose again.

**Figure 5:**
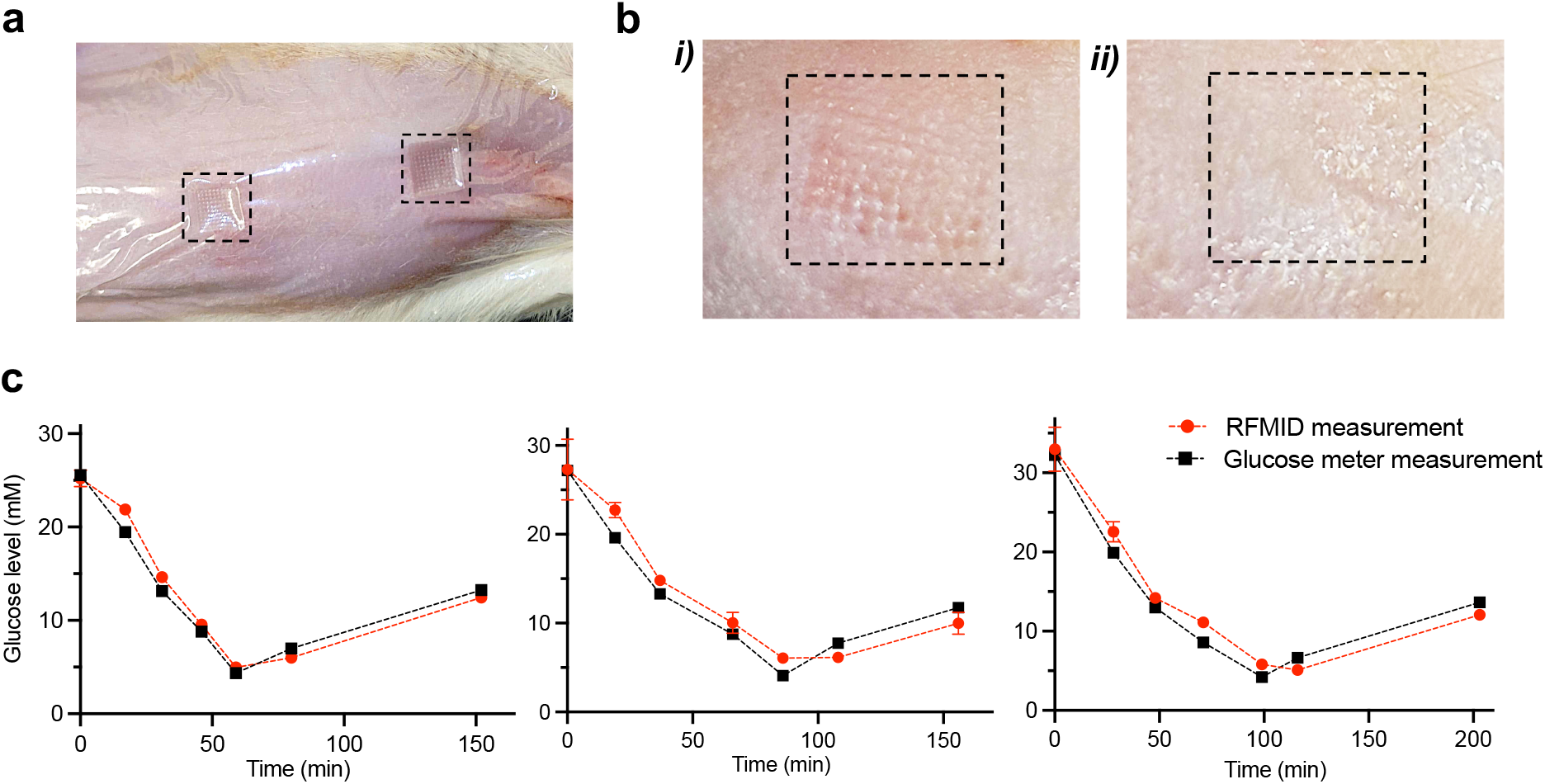
*In vivo* glucose detection in diabetic rats. **a**, RFMID patches were applied into the dorsal skin of awake rats and fixed with Tegaderm tapes for 5 min. **b**, Magnified images of the trace of a patch on the skin (***i***). The skin is recovered after 15 min (***ii***). **c**, RFMID measurement of glucose levels in three different diabetic rats over 150-200 mins. The rats were injected with 4 IU kg^−1^ dose of human recombinant insulin (at t = 5 min; after baseline measurement) subcutaneously or 30% glucose solution (at t = 63, 93, 109 min for different rats and after reaching hypoglycemia condition) intraperitoneally to reduce or increase the blood glucose level, respectively. Six ranges of glucose levels were targeted in total: T0: 25 – 35 mM, T1: 15 – 25 mM, T2: 10 – 15 mM, T3: 5 – 10 mM, T4: 3 – 10 mM, and T5 > 10 mM. After reaching each range, three RFMID patches were applied to the dorsal skin of rats for 5 mins. For each timepoint, we collected blood samples from the tail vein before and after applying RFMID patches, measured the glucose levels using a hand-held glucose meter, and reported the average to compare with RFMID measurement. These results correlated closely with a time lag of 3-14 mins and highlight the inter-individual variability in the insulin or glucose response.

The RFMID patches for glucose detection were applied at different time points and kept on the skin for 5 min (**Figure 5c & S8**). These results demonstrate that the RFMID device can track the falling and rising glucose concentrations in animal models. In parallel, we compared the RFMID results with conventional glucose measurements from a handheld glucose meter using blood samples collected from the rat tail vein and observed that both sets of results correlated well with a time lag of 3-14 mins. This time lag is attributed to the locations of glucose and differing transport efficiencies between ISF and circulating blood. Patients with diabetes also showed a time lag of 4–10 min in the change of ISF glucose levels relative to blood glucose concentrations^34^. The differences between glucose responses in individual rats clearly show the inter-individual variability, even under controlled conditions with genetically similar animals. This has important implications for human patients, who are genetically diverse and exposed to different environmental conditions. Thus, these results highlight the necessity of personalized glucose monitoring using a simple and reagentless approach.

### Outlook

Herein, we demonstrated the first technology to combine HMN arrays with aptamer probes to summon their merits for reagentless and minimally invasive target detection. We showed a comprehensive characterization of our HMN patches functionalized with aptamer probes where addition of the aptamer probes did not have a significant effect on the swelling ability or mechanical strength of the patches. Experiments in skin specimens equilibrated with varying concentrations of glucose or ATP indicate that our sensor has high sensitivity and specificity to detect clinically relevant concentrations of both analytes, with a LOD of 1.1 mM for glucose and 0.1 mM for ATP. The RFMID assay’s response time to capture and detect the target analytes is only 2 min because of the swelling capabilities of MeHA that enable rapid ISF access and the reagentless detection mechanism. This fast response time is an exceptional improvement over previously reported MN-based diagnostic devices (**Table S1**). *In vivo* experiments in awake diabetic rats confirmed the RFMID ability to measure changes in glucose with no need for adding any reagents and highlighted the RFMID platform’s capacity to detect inter-individual variations in glucose response between animals - a critical feature for clinical implementation. Importantly, RFMID measurements closely matched those obtained with standard clinical glucose meters.

Although it is beyond the scope of this work, we believe our system could be modified for continuous, real-time measurement in a minimally invasive manner. Aptamer probes have been recently employed to the continuously measure biomolecules *in vivo* using electrochemical sensors where the structure switching characteristics of aptamers were integrated with a redox reporter to produce a concurrent electrochemical signal^35^. The aptamer-based electrochemical detection has been applied for continuous measurement of different metabolites and drugs, however, the complicated and invasive design (i.e., insertion into the vein that requires surgery) and/or the short monitoring capacity (a few hours) are key shortcomings^35–37^. We envision modifying our HMN-based biosensor integrated with the aptamer probes for continuous measurement via tracking electrochemical signals. Important considerations to that end would be to develop a conductive HMN patch that can be employed as the working electrode, as well as fabricating MN-based reference and counter electrodes. With such modifications, an integrated technology to continuously collect individual patient molecular profiles in a minimally invasive manner can be deployed, allowing continuous and prolonged measurement of any targets of interest, such as drugs with narrow therapeutic range.

Finally, we emphasize that the RFMID system is a platform that could be readily modified to measure other circulating analytes *in vivo*, for which aptamer probes are available, thus making it potentially a versatile tool for diverse biomedical applications.

## Methods

### Materials

The Pharma-grade sodium Hyaluronic acid (HA, MW 300KDA) was purchased from Bloomage Co., Ltd (China). 1 × PBS, Dimethyl sulfoxide (DMSO, 25-950-CQC), was purchased from Corning, USA. Irgacure 2959 (photo initiator, PI), N’-methylenebisacrylamide (MBA), methacrylic anhydride (MA), glucose and ATP solution and other chemicals were purchased from Sigma Aldrich (Canada). 100 mM ATP solution (R0441) was purchased from Thermo Fisher. The porcine ear skin was obtained from a local supermarket. All the aptamer and displacement strand were purchased from Integrated DNA technologies. Sequence of ATP^30^ and glucose^29^ aptamer and competitor strands were obtained from literature. Sequence and modification of all aptamer and competitor strands are indicated in Table 1.

**Table 1.**
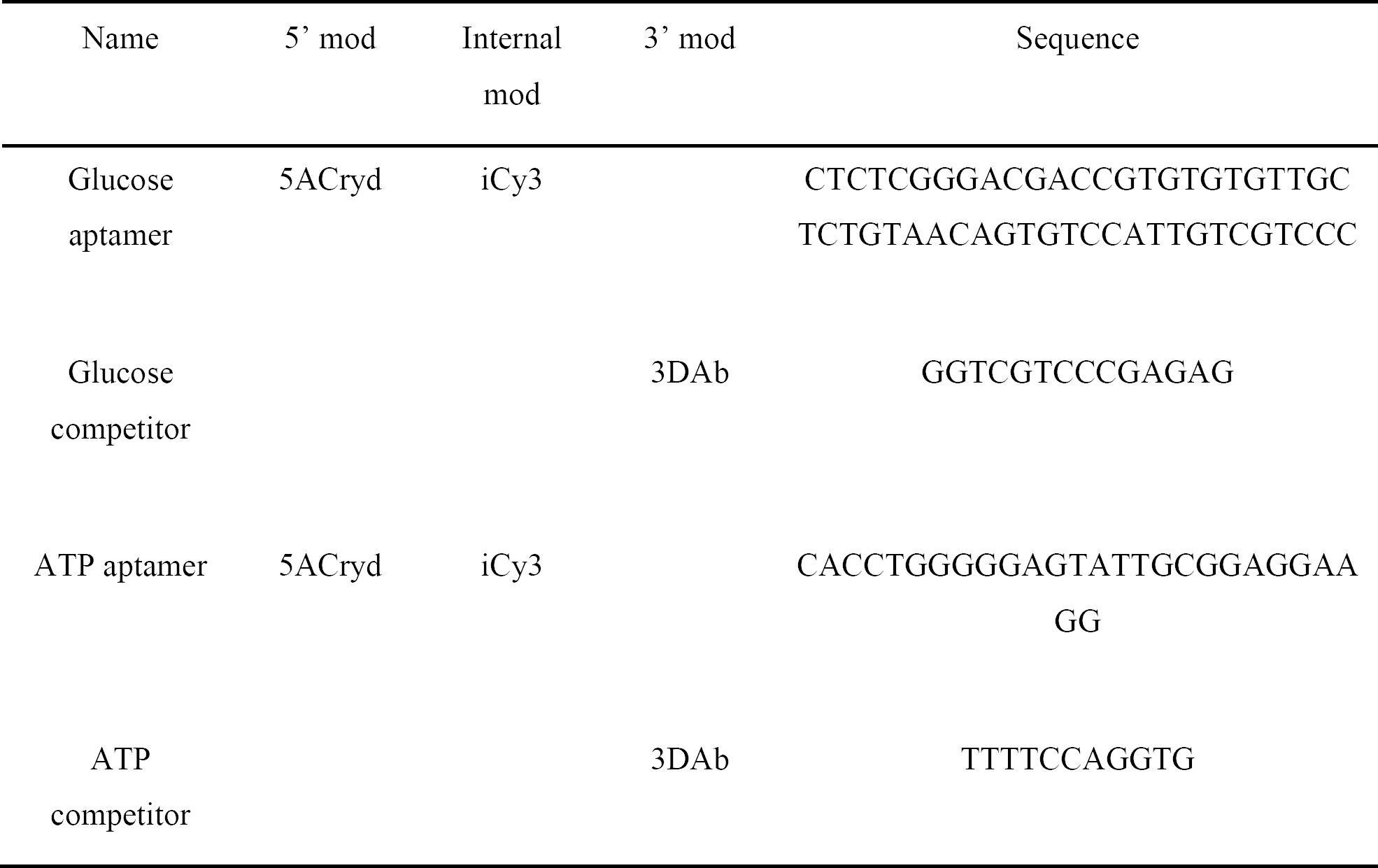
Sequence and modification of aptamer and competitor strands.

### Synthesis and characterization of MeHA

MeHA was synthesized based on the modified protocol established by Poldervaart et al^22^. 2.0g HA was dissolved in 100 mL Millipore water and stirred overnight under 4 degree for complete dissolving. Subsequently, 1.6 mL MA was added into HA solution and 3.6 mL of 5N NaOH solution was added to adjust the solution to pH 8-9. The mixture was stirred overnight under 4 degree to complete the reaction. Next, MeHA was precipitated by acetone and washed three times with ethanol. Subsequently, precipitated MeHA was redissolved in Millipore water and was dialyzed for 2 days to remove the impurity. The purified MeHA was lyophilized for 3 days. Eventually, 2 -5 mg of MeHA was dissolved in 1 mL Deuterium oxide (D_2_O) (Sigma Aldrich, 151882) and then tested with 300MHz ^1^HNMR with 10 ms time scale. The degree of methacrylate modification was determined by integration of methacrylate proton signals at 6.1, 5.7, and 1.8 ppm to the peak at 1.9 ppm related to the N-acetyl glucosamine of HA^23^.

### Fabrication of RMFID

For each RMFID patch, 50 mg MeHA, 1 mg PI and 1 mg of MBA were dissolved in 1.25 mL of glucose or ATP aptamer binding buffer. The MeHA solution was then sonicated for 5 min to remove the bubbles. Subsequently, 0.5mL of MeHA solution was deposited on a negative polydimethylsiloxane (PDMS) mold (Micropoint, Singapore), and degassed for 90s. After drying at room temperature for 5 hours, another 0.75 mL of MeHA solution was casted on the mold followed by drying at room temperature for 10 hours. Next, 10 μL of aptamer-quencher strand solution composed of 1μM aptamer and 10 μM corresponding quencher strand with 15 min pre-hybridization was loaded on the HMN followed by drying at 45 degrees for 30 min. Dry HMN patches were then crosslinked by UV light with 360 nm wavelength for 15 min. The RMFID patches were washed twice with 10 μM of glucose or ATP aptamer binding buffer and dried under 45 degrees. Last, MN patches were carefully separated from PDMS molds and further crosslinked for 5 mins. After being trimmed, the RMFID patches were observed under a fluorescence microscope (Nikon, Ti2). The fabricated needles of the RMFID patch were 850 μm in height, 250 μm in base width and 500 in internal spacing.

### Chemical characterization

The Fourier-transform infrared spectroscopy (FTIR, Bruker Hyperion 3000 FTIR Microscope) was conducted to study the crosslinked degree of HMNs and aptamer functionalization efficiency. The following four samples were made in 1 mL DI water: two samples of 50 mg/mL MeHA solution containing 1 mg/mL photo initiator (MeHA); one sample of 50 mg/mL MeHA solution containing 1 mg/mL photo initiator and 1 mg/mL MBA (MeHA+MBA); and one sample of 50 mg/mL MeHA solution containing 1 mg/mL photo initiator and 1 μM aptamer (MeHA+APT). One sample of MeHA, MeHA+MBA and MeHA+APT were crosslinked under UV exposure for 20 min, and then their FTIR spectrum is recorded from 4000 to 400 cm^-1^ and were compared with not-crosslinked MeHA sample.

### Mechanical test and skin penetration efficiency

The RMFID patches were applied on rat dorsal skin or on the porcine skin for 15 min. Subsequently, the trace on skin was recorded by digital camera every 5 min for 15 mins. The mechanical strength of MN patches was measured using Instron 5548 micro tester equipped with 500N compression loading cell. For each test, the HMN patch was placed flat on its backside (tips facing upwards) on a compression platen. The distance between two platens was set to 1.5 mm. A vertical force was applied (at a constant speed of 0.5 mm/min) by the other platen. The compression loading cell capacity was set to 70 N. The load (force; N) and displacement (distance; mm) was recorded by the testing machine every 0.1 s to create the load-displacement curve.

### *In vitro* cytotoxicity assay (evaluation of biocompatibility)

The biocompatibility of RMFID was investigated using mouse fibroblast cells (NIH-3T3). Cells were seeded at a density of 50,000 cells per well in a 96-well plate with a final volume of 100 μL. Subsequently, cells were exposed to 10 μL of sample solution for 24 hours. The 5mL sample solution contains 50 mg MeHA, 1mg MBA, 1mg photo initiator and 10 μL of 1 μM ATP or glucose aptamer solution. 10 μL of DMEM medium solution was used as control. After sample exposure, 10 μL of the 5 mg/ml Methylthiazolyldiphenyl-tetrazolium bromide (MTT) (Sigma Aldrich, M5655) solution was added to all wells. Next, the plate was incubated in the absence of light for 3 hours. 150 μl of DMSO was added and gently pipetted to all wells to break up cells and release the formazan crystals. The absorbance of the samples was then obtained at 540 nm using a spectrophotometer.

### Swelling studies

A 1.4wt % agarose (Sigma Aldrich, A0169) hydrogel was prepared in DI water. The dry mass (W_0_) of HMNs were measured before applying through agarose. Then the HMNs were penetrated to the agarose through a layer of parafilm and swelled for 10 min. Next, wet mass (W_t_) of the swelled HMN patches was measured. The swelling ratio of HMNs was calculated based on the below formula:

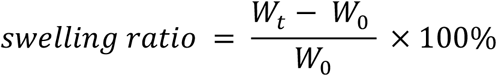

### Assessing Rhodamine B recovery rate

A 1.4wt% agarose hydrogel containing 100 mg/mL (C_0_) Rhodamine B (Rho B) (Sigma Aldrich, R6626) was prepared. HMN patches with crosslink time of 5, 10, 15, 20 min were weighted, and their dry mass (W_0_) was recorded. Next, HMNs were punched into RhoB agarose through parafilm for 10 min. After measuring the wet mass (W_t_), the swelled HMN was mixed with 300 μL (V) Millipore water in a centrifuge tube followed by a 5 min centrifugation at 10K rpm^4^. Subsequently, 80 μL of recovered solution was transferred into a 96-well plate to measure the absorbance at 552 nm. The Rho B recovery rate were calculated based on the following formula.

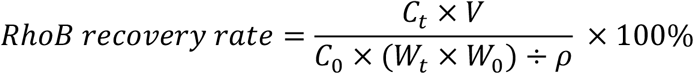

In the equation, C_0_ refers to the initial RhoB concentration (100 mg/mL), C_t_ is the detected RhoB concentration recovered from MN, V is the volume of recovered solution (300 μL), and (*W*_*t*_ − *W*_0_) ÷ *ρ* is the volume of solution absorbed by MN.

### Assessing glucose recovery rate

To evaluate the capability of the RFMID for glucose recovery, three groups of samples were prepared: a group of blank HMN patches without aptamer probe functionalization and two groups of HMN patches functionalized with glucose aptamer probes (Glu apt HMN). After measuring the dry mass (W_0_), HMNs were penetrated to 1.4 wt% agarose containing varying glucose concentration of 3.5, 5, 10, 20 mM for 10 min. After measuring the wet mass (W_t_), one group of HMN patches functionalized with aptamer probes was sonicated for 10 min and named as Glu apt-US HMN. Subsequently, all the HMN were mixed with 300 μL (V) Millipore water in a centrifuge tube followed by 5 min centrifugation at 2,100 rcf. Subsequently, 250 μL of recovered solution was transferred into a 96-well plate and the recovered glucose concentration was measured using a glucose (GO) assay kit purchased from Sigma (GAGO20). Glucose recovery rate was defined by the following formula.

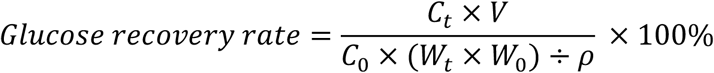

C_0_ refers to the initial glucose concentrations (3.5, 5, 10, 20 mM), C_t_ is the detected glucose concentration recovered from MN, V is the volume of recovered solution (300 μL), (*W*_*t*_ − *W*_0_) ÷ *ρ* is the volume of solution absorbed by HMN.

### *In vitro* glucose and ATP measurement

The Fluorescence intensity (FI) of the RFMID was recorded by the fluorescent microscope from the base side. The RFMID patches with ATP or glucose aptamer probe were applied on 1.4 wt% agarose hydrogels containing various concentrations of ATP (0, 0.25, 0.5, 1, 2, 4 mM) or glucose (0, 3.5, 5, 10, 20, 32 mM) for 10 min, respectively. Next, the fluorescent intensity of the RFMID after target detection was recorded. Finally, the corresponding needles were identified and the fluorescence intensity difference before and after target capturing were measured and calculated by subtracting FI before target capture from the FI after target capture.

### Assessing specificity of RMFID

The RMFID functionalized with glucose or ATP aptamer probe was punched into the 1.4 wt% agarose containing 10 mM glucose and 0, 1, 2, 4 mM ATP, or 1mM ATP and 0, 5, 10, 20 mM glucose for 10 min, respectively. The fluorescence intensity of RMFID before and after being applied on agarose was recorded by fluorescent microscope.

### *Ex vivo* glucose and ATP measurement

After being rinsed with DI water and trimmed to 1 cm by 1 cm square, porcine ear skins were equilibrated in 1XPBS with various concentrations of ATP (0, 0.25, 0.5, 1, 2, 4 mM) or glucose (0, 3.5, 5, 10, 20, 32 mM) overnight. Subsequently, fluorescent intensity RFMID patches were recorded with the fluorescence microscope from the base side. Next, ATP or glucose RFMID patches were applied on porcine skin equilibrated with ATP or glucose for 5 min, respectively. Tegaderm tape (3M) was used to fix RFMID patches on the skin. After drying under room temperature, the RFMID patches were observed under the fluorescence microscope and their fluorescent intensity was recorded. Similar to the *in vitro* experiment, the fluorescent intensity difference of RFMID before and after target capturing was calculated.

### *In vivo* glucose measurement in diabetic rats

Animal studies were performed in accordance with the Guidelines for the Care and Use of Laboratory Animals and the Animal Welfare Act Regulations; all protocols were approved by the University of Toronto Institutional Animal Care and Use Committee. An established model of T1D, the streptozotocin (STZ)-induced diabetic rat, was used to explore the *in vivo* performance of RFMID. Male Sprague Dawley rats (Charles River, 100-150 gr) were injected with STZ (65 mg/kg i.p.) that destroys the host’s pancreatic beta-cells secreting insulin^38^. After the STZ injection, the rats were monitored for 1 week and their blood glucose was measured every 2 days using a glucose meter (OneTouch® Ultra®, LifeScan, Inc., USA). Diabetic rats with blood sugar stabilized above 17 mM were selected for this study. Before starting experiments, the rats were fasted for 5 hours. Rat skin was then shaven, treated with hair removal cream, and dried prior to MN patch application. RFMID patches for ISF glucose detection were prepared and their fluorescence intensity was measured. The baseline blood and ISF glucose level of rats were measured by glucometer and RFMID, respectively. Subsequently, 4 units of insulin were injected to the rats subcutaneously and blood glucose levels were tracked by glucometer every 5 min. The RFMID patches were applied on rats’ skin and fixed with Tegaderm tape for 5 min, when blood glucose level decreased to certain ranges (5 ranges in total): T0: 25 – 35 mM, T1: 15 – 25 mM, T2: 10 – 15 mM, T3: 5 – 10 mM, T4: 3 – 5 mM. After reaching to hypoglycemia regime, 0.5 mL of 30% glucose solution was intraperitoneally injected into the rats, followed by another two time-point ISF glucose measurements by RFMID. After drying at room temperature, RFMID patches were observed under the microscope and their fluorescence intensity was recorded. Rats’ ISF glucose levels were calculated by interpolating the fluorescence intensity difference before and after detection into the *ex vivo* RFMID glucose detection calibration curve.

### Statistical analysis

All the statistical analysis was conducted using GraphPad Prism 9. The statistical difference between groups in biocompatibility test, RFMID specificity test and swelling experiment was analyzed using ordinary one-way ANOVA with Tukey’s multiple comparison test. In glucose recovery experiment, the statistical difference between different HMN groups and various glucose concentrations were analyzed using two-way ANOVA with Geisser-Greenhouse correction. The significance of statical difference was calculated with 95% confidence interval (P<0.05) and shown in GP style (0.1234 (ns), 0.0332 (*), 0.0021 (**), 0.0002 (***), <0.0001 (****)) in graphs. Each experiment contains three parallel replicates. All data is expressed as mean ± s.d. For *in vitro* and *ex vivo* glucose and ATP measurement, fluorescence signal raw data after subtracting the background signal was used to calculate the calibration curves using the sigmoidal 4PL nonlinear regression model. The LOD is defined as the minimum target concentration that can be detected by our RFMID device and calculated by interpolating the mean fluorescence signal of control group plus three times of s.d. into the corresponding calibration curve^39^.

## Data availability

The data that support the findings of this study are available from the corresponding author upon reasonable request.

## Acknowledgment

This research was supported by the Natural Sciences and Engineering Research Council of Canada, Discovery grant, Poudineh’s University of Waterloo start-up funding and Center for Bioengineering and Biotechnology Seed funding. We thank Karan Dhingra and Sarah Odinotski for their help on preparing the RFMID patches. We thank Dr. Nafiseh Moghimi with her help for performing the biocompatibility experiment. The authors acknowledge Waterloo Advanced Technology Laboratory, Nuclear Magnetic Resonance facility, Center for Advanced Materials Joining, and Giga-to-Nanoelectronic center for their assistance with FTIR, HNMR, mechanical strength, and SEM measurement.

## Author contributions

H.Z., A.Gh., and M.P. conceived the initial concept and designed experiments. H.Z., A.Gh, P.Gh and M.S. executed experiments and analyzed the data. H.Z., A.Gh., and M.P. wrote the manuscript. H.Z., A.Gh, P.Gh, M.S., A.G., and M.P. authors edited, discussed, and approved the whole paper.

## Additional information

Online Supplementary Information accompanies this paper.

---

**Table 1.** Comparison of RFMID’s response time with similar MN-based sensors

|  | Capture | Washing | Blocking | Detection | Washing | Total time |
| --- | --- | --- | --- | --- | --- | --- |
| **FLISA<sup>1</sup>** | 30 s-20 min | 10 min | 30 min | 120 min incubation with Ab + 30 min incubation with plasmonic fluor | 10 min | 200 min-220 min |
| **Aptamer decorated MN array<sup>2</sup>** | 60 min | 10 min | - | 60 min incubation with FAM labeled aptamer | 10 min | 140 min |
| **Encoded MNS<sup>3</sup>** | 40 min | 10 min | - | 120 min incubation with detection Ab | 10 min | 180 min |
| **RFMID** | 2 min | - | - | - | - | 2 min |

**Figure S1.**
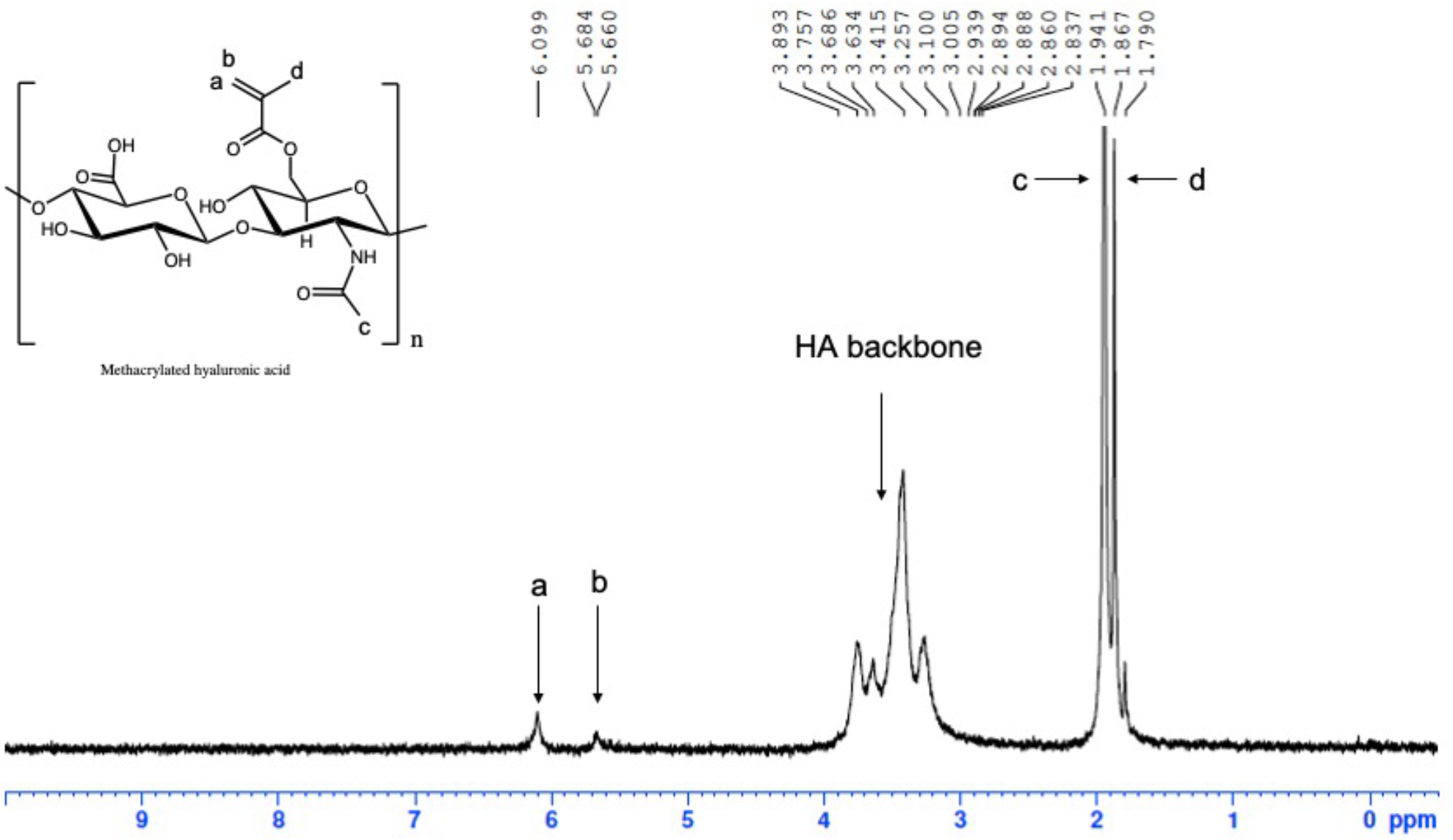
^1^H NMR spectra of MeHA. MeHA was characterized with 300MHz ^1^HNMR with 10ms time scale to determine the degree of methacrylate modification.

**Figure S2.**
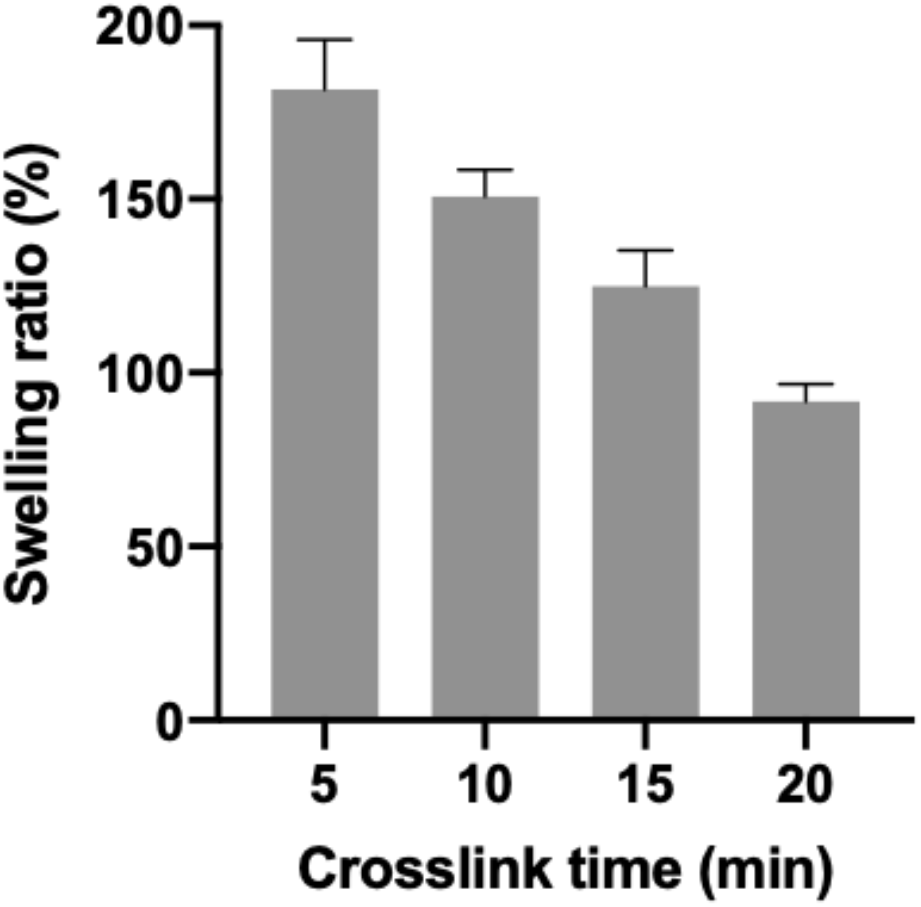
Characterization of crosslinked MeHA-HMN. MeHA-HMNs with various crosslinking time (5, 10, 15, 20 min) were characterized based on their swelling ability.

**Figure S3.**
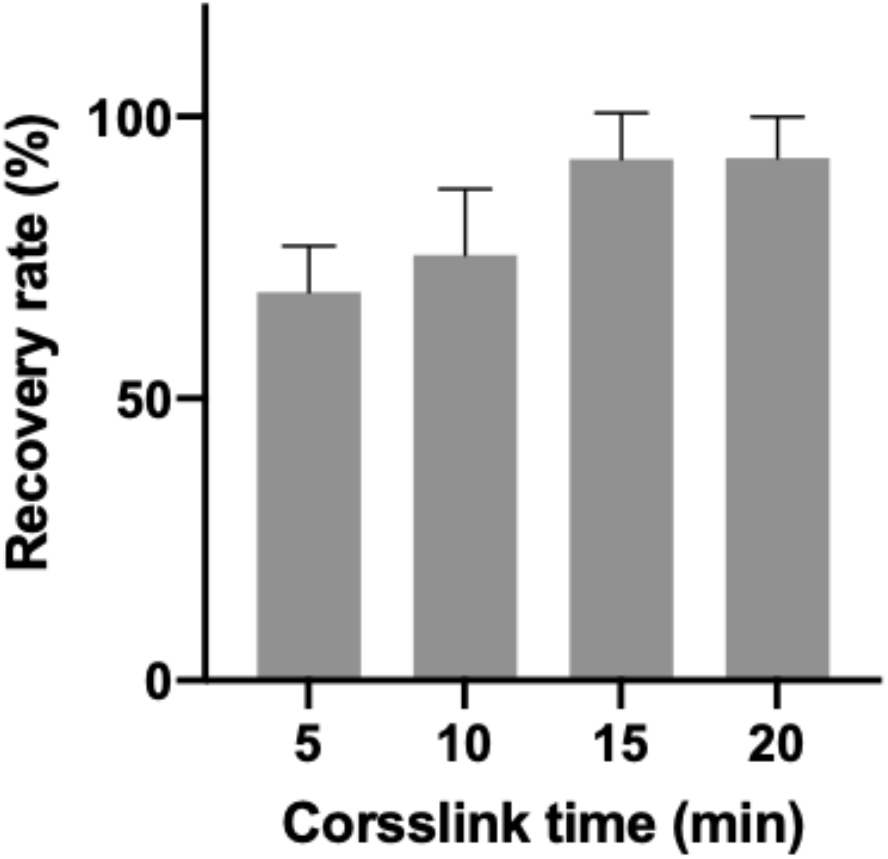
Characterization of crosslinked MeHA-HMN. MeHA-HMNs with various crosslinking time (5, 10, 15, 20 min) were characterized based on their Rhodamine B recovery rate.

**Figure S4.**
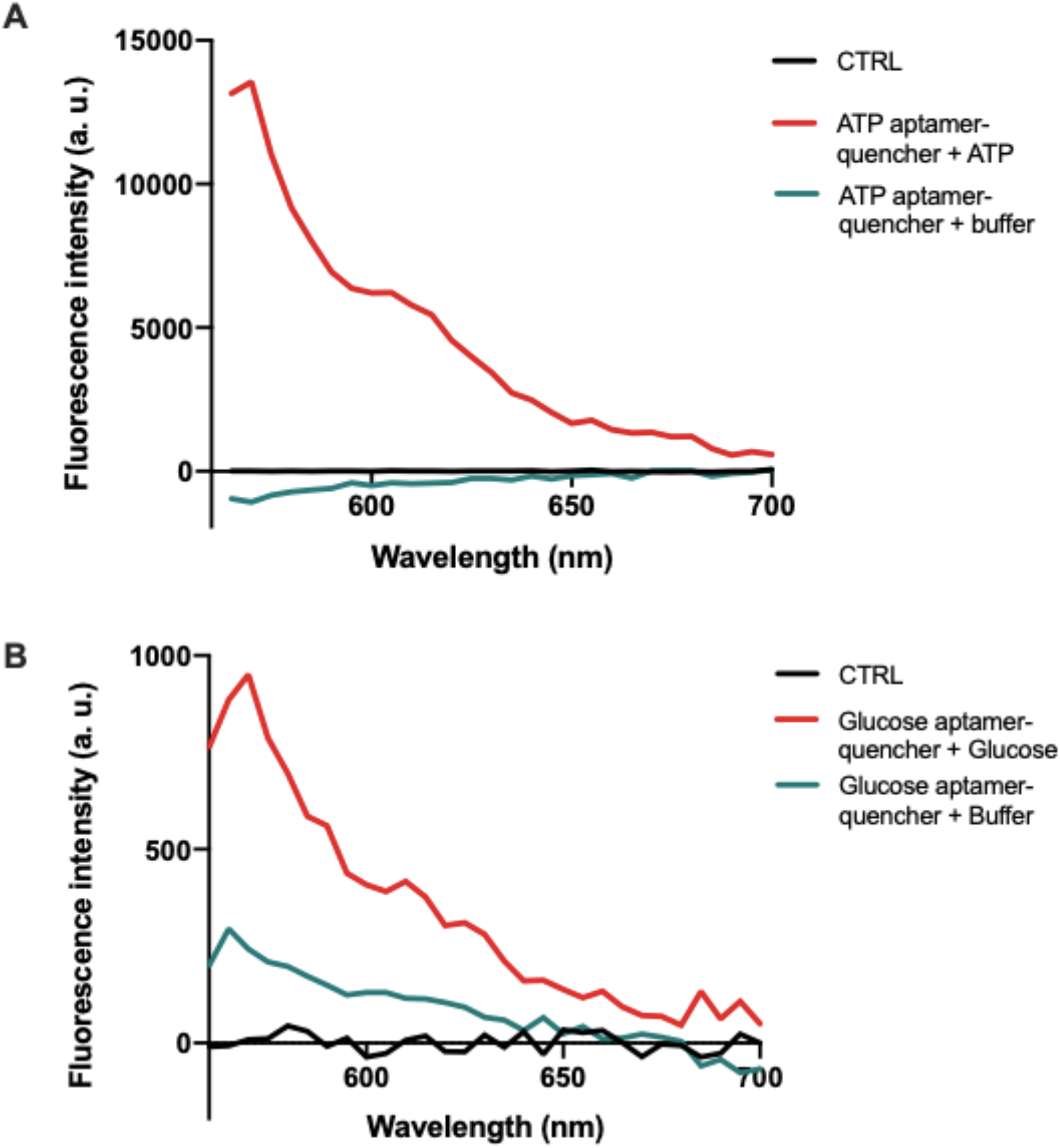
Assessment of aptamer binding efficiency. The target binding efficiency of **A)** ATP and **B)** glucose aptamer was tested with spectrophotometer. A spectral scanning measurement was conducted with the excitation of 530 nm, and an emission from 550 nm to 700 nm. Buffer solution was used as background control (CTRL). The aptamer probe was hybridized to its corresponding quencher for 30 min before adding the targets. Subsequently, the fluorescence intensity difference was measured after capturing target for 30 min.

**Figure S5.**
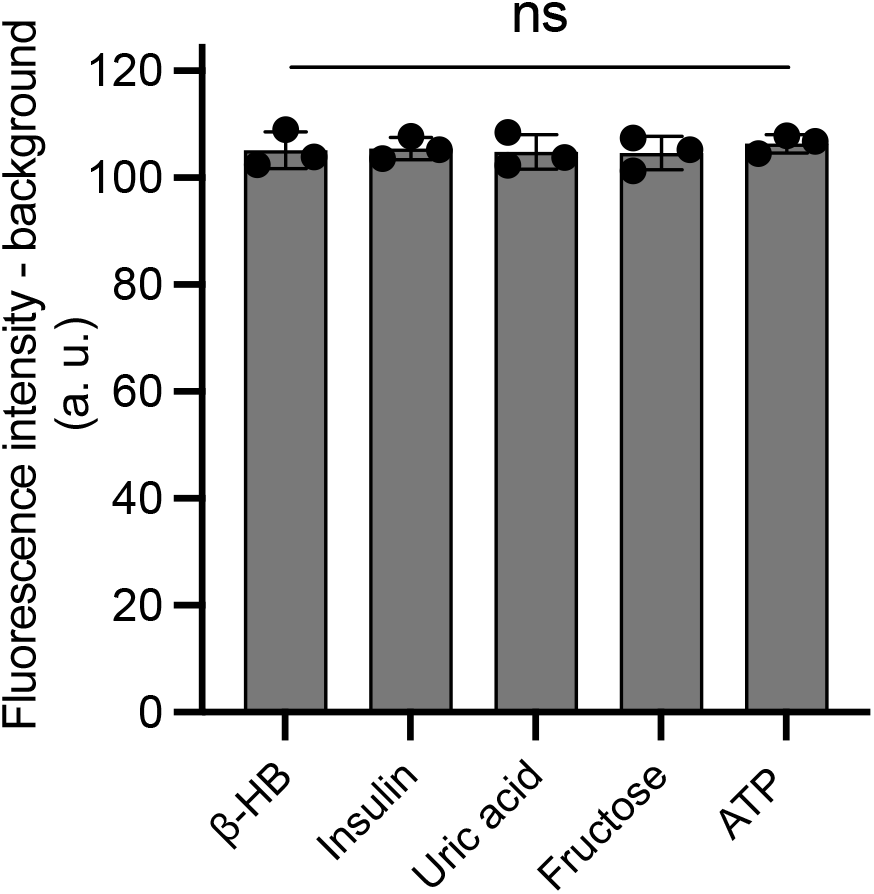
The cross-reactivity of RFMID device for glucose capture in the presence of common interfering agents was studied. In this experiment, agarose hydrogel was loaded with a glucose concentration of 20 mM and β-HB concentration of 10 mM or insulin concentration of 10 nM or fructose concentration of 0.5 mM or uric acid concentration of 0.5 mM or ATP concentration of 4 mM. Data is expressed as mean ± s.d. n = 3 replications per group (ns, not significant).

**Figure S6.**
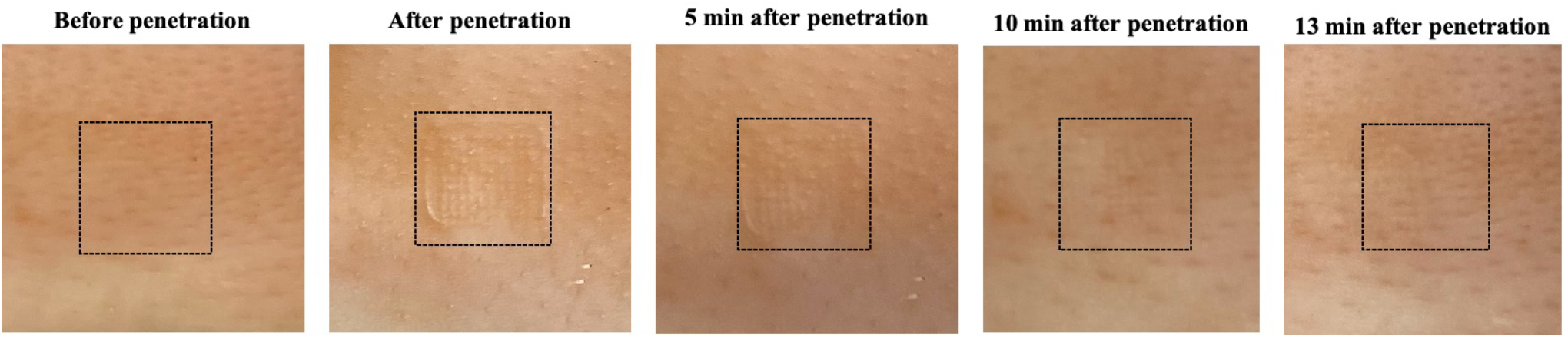
*Ex vivo* porcine skin penetration efficiency test using RFMID. Glucose RFMID patches were applied to the porcine ear skin for 5 min with index finger pressure. After removing the RFMID patches, skin samples were imaged with digital camera for 13 min with a 5 min interval to observe the micro-sized holes left by RFMID.

**Figure S7.**
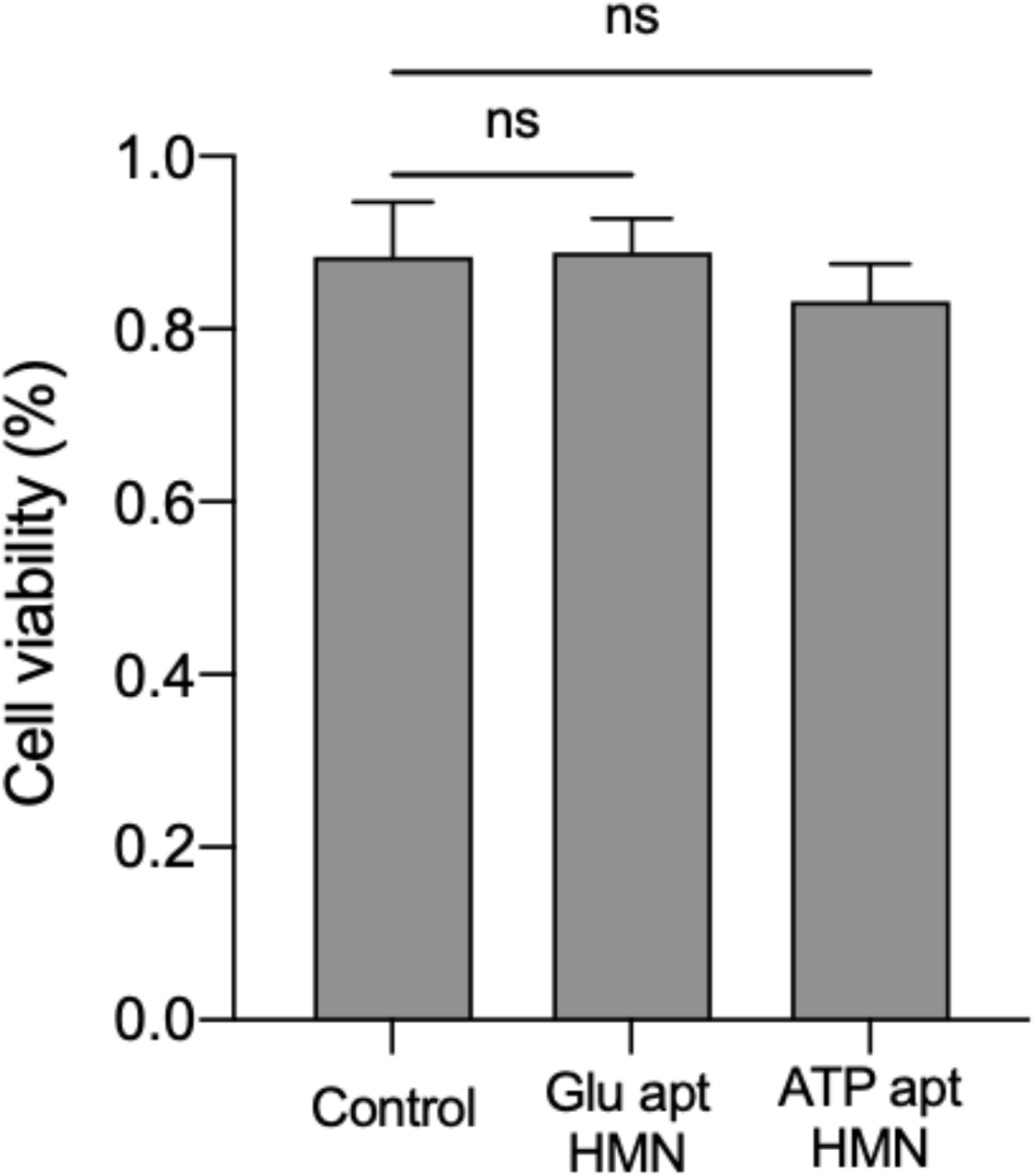
Biocompatibility test of aptamer MeHA HMN. Mouse fibroblast cells were cultured at 100,000 cells per well in a 96-well plate and exposed to 10 μL of MeHA samples solution for 24 hours. Sample solution composed of 50 mg MeHA, 1mg MBA, 1mg photo initiator and 10 μL of 1 μM ATP or glucose aptamer solution. Cells in control group were exposed to cell culture media. Each group has three replicates. Subsequently, cell viability was detected with MTT assay.

**Figure S8.**
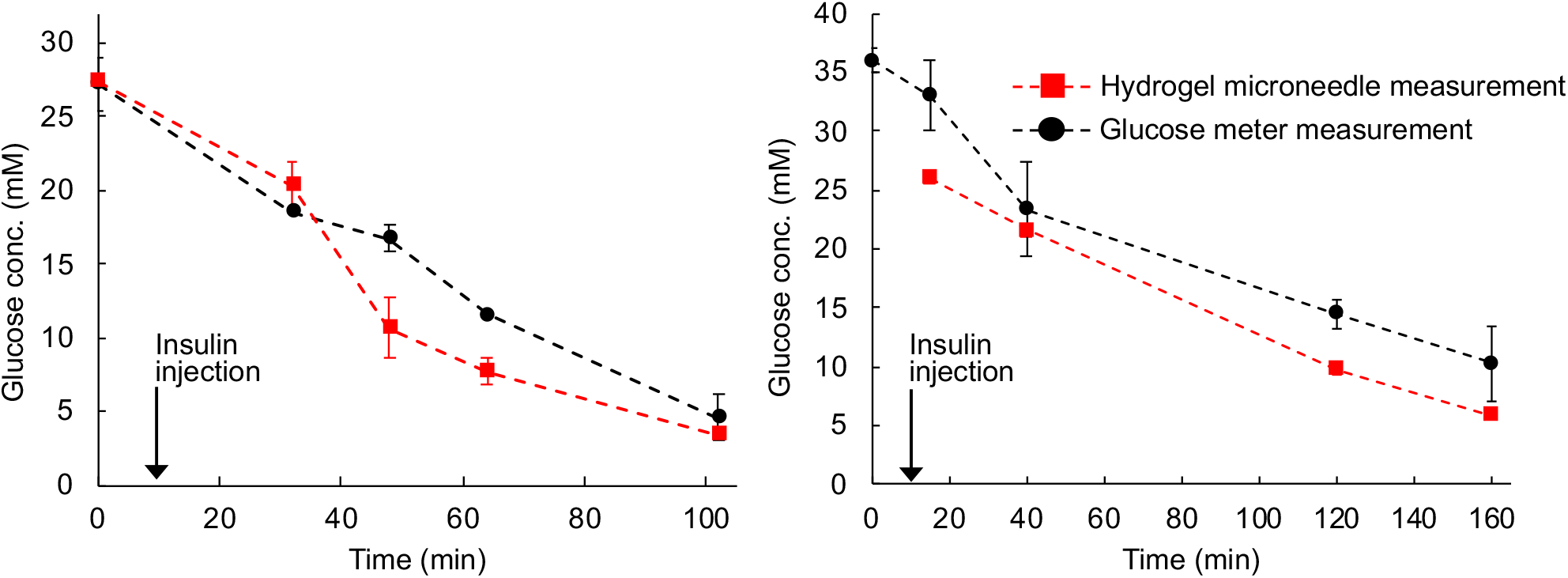
Glucose measurement in live diabetic animals using RFMID. Insulin bolus was injected after the baseline measurement which results in decrease in the glucose level to hypoglycemia regime.

